# Distinct Patterns of Adaptive Decision-Making in Mouse Models of Neurodevelopmental Disorders

**DOI:** 10.64898/2026.10.14.759086

**Authors:** James Hilber, Kerda Pulk, Takumi Nakamura, Kurara Honda, Tadafumi Kato, Atsushi Takata, Takeru Matsuda, Aya Ito-Ishida

## Abstract

Neurodevelopmental disorders (NDDs) arise from diverse genetic alterations that can affect adaptive decision-making, but quantifying specific components of cognitive processes remains challenging. We developed a sequence of deterministic and probabilistic reversal learning tasks using an automated operant platform and applied reinforcement-learning models to quantify latent cognitive processes in mice. We applied this framework to three genetic NDD models: *Mecp2*^+/–^ (Rett syndrome), *MECP2*-Tg1 (*MECP2* duplication syndrome), and *Kmt2c*^+/fs^ (*KMT2C* haploinsufficiency) mice. *Mecp2*^+/−^ mice showed heterogeneous impairment in acquiring deterministic reversal learning, accompanied by reduced task engagement and increased premature responses. *MECP2*-Tg1 mice exhibited mild impairment of deterministic reversal learning but preserved performance in probabilistic reversal learning. Although *Kmt2c*^+/fs^ mice showed largely preserved behavioral performance, reinforcement-learning modeling revealed increased learning from unrewarded outcomes and faster forgetting of unchosen options, which were not apparent from overall performance. These findings demonstrate that combining quantitative reversal learning tasks with computational modeling can distinguish specific alterations in cognitive processes, providing an effective framework for characterizing cognitive dysfunction across mouse models of NDDs.

## Introduction

Neurodevelopmental disorders (NDDs) are associated with diverse clinical manifestations, reflecting, in part, their genetic heterogeneity^1,2^. Cognitive impairments in NDDs span multiple domains, including learning and memory, attention, and cognitive flexibility, and can lead to impaired adaptive decision-making essential for everyday life^3,4^. However, how diverse genetic alterations give rise to distinct cognitive phenotypes remains poorly understood. Addressing this question requires behavioral approaches that enable quantitative assessment of specific cognitive processes in both humans and animal models. Such approaches may help characterize both shared and distinct patterns of cognitive dysfunction across genetically diverse NDDs.

Adaptive decision-making involves adjusting choices as circumstances change and has been widely studied using reversal learning tasks across species^5–7^. In these tasks, individuals choose among multiple options and receive rewards, with outcome contingencies changing over time. In deterministic reversal learning tasks, choices are consistently rewarded or unrewarded, providing clear feedback. In contrast, in probabilistic reversal learning tasks, outcomes are uncertain, requiring individuals to integrate experience across multiple trials to adjust their choices^5,8^. Multiple factors, including engagement, motivation, impulsivity, and sensitivity to feedback, can influence performance in these tasks^9,10^. In addition, latent mental processes can also affect performance. These processes are not readily apparent in behavioral performance but can be inferred by using computational modeling^11^. For example, reinforcement-learning models quantify latent properties of adaptive decision-making, such as how much choices are updated after rewarded versus unrewarded outcomes and how quickly the values of unchosen options are forgotten^12^.

Previous studies have examined reversal learning in mouse models of NDDs to understand the biological basis of cognitive alterations^13–15^. These studies have revealed diverse phenotypes, ranging from impaired cognitive flexibility to alterations in specific aspects of decision-making. Combining behavioral tasks with computational modeling has further revealed alterations in processes such as value updating, as demonstrated in *Nrxn1*α-deficient mice^16^. However, most studies have focused on individual genetic models, and direct comparison across models remain difficult because of differences in task structure, reward contingencies, and analytical approaches. Applying a shared behavioral and computational framework to multiple genetic models may therefore provide insight into the diversity of decision-making phenotypes across NDD models.

In this study, we developed a sequence of reversal learning tasks to quantify adaptive decision-making in mice. We conducted these tasks using an automated home cage system, Operant House, in which mice performed tasks during the night without experimenter handling^17^. This approach allowed us to systematically measure multiple aspects of behavior, including task engagement, cognitive flexibility, and premature responses, under minimally stressful conditions. We further applied computational reinforcement-learning models to characterize processes underlying decision-making. To examine genetically distinct NDD models within the shared framework, we tested *Mecp2*^+/−^ mice, which model *MECP2* loss of function and reproduce key features of Rett syndrome^18^; *MECP2*-Tg1 mice, which model *MECP2* overexpression associated with *MECP2* duplication syndrome^19^; and *Kmt2c*^+/fs^ mice, which model *KMT2C* haploinsufficiency and exhibit autism-related behavioral phenotypes^20^. Using the shared behavioral and computational framework across these models, we identified distinct behavioral phenotypes and computational alterations that were not readily apparent from overall task performance.

## Results

### Development of a behavioral task sequence to measure adaptive decision-making in mice

To examine adaptive decision-making in mouse models, we developed a sequence of touchscreen operant conditioning tasks using Operant House (**Figure 1A, B**)^17^. The task sequence consisted of a visual discrimination (VD) task to establish an association between panel touches and water reward, followed by a deterministic reversal learning (DRL) task with 100–0 reward contingencies and three probabilistic reversal learning (PRL) tasks with progressively increasing reward uncertainty (80–20, 70–30, and 60–40; **Figure 1B**). The VD and DRL tasks were repeated across days until mice reached predefined criteria before progressing to the subsequent tasks. The criterion for the VD task was a correct rate above 65% for 2 consecutive days, whereas the criterion for the DRL task was a correct rate above 75% for 1 day or above 70% for 2 consecutive days.

**Figure 1.**
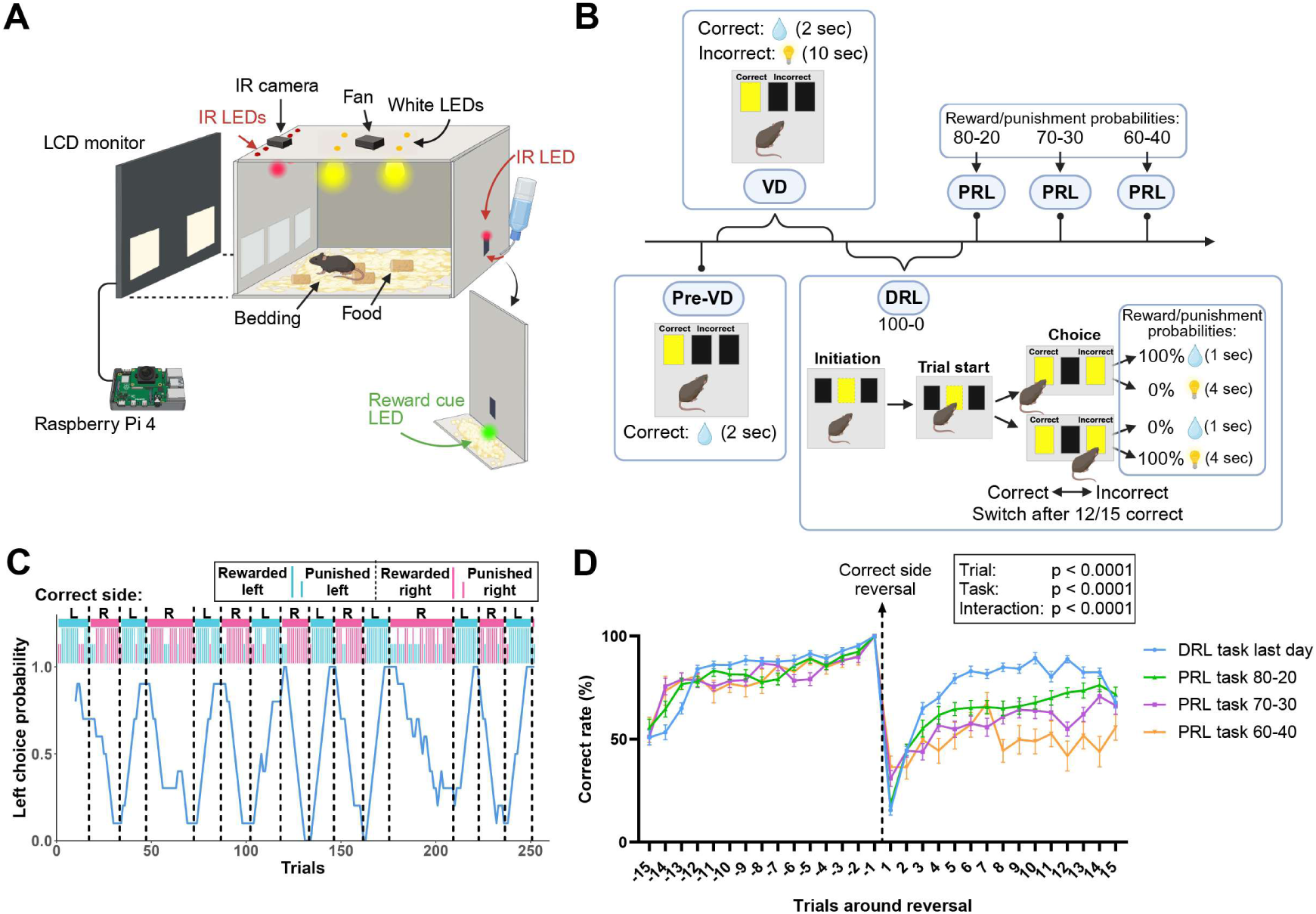
Design of the Operant House, task sequence structure, and an example of task performance in wild-type male mice. (A) Schematic of Operant House. (B) Task sequence consisting of pre-visual discrimination (Pre-VD), visual discrimination (VD), deterministic reversal learning (DRL), and probabilistic reversal learning (PRL) tasks. (C) Representative choice behavior of a WT mouse during the last day of the DRL task. Blue and pink shading indicates periods in which the left and right panels, respectively, were designated as the correct choice. Long ticks represent rewarded trials. Short ticks represent punished trials. Left choice probability for each trial was calculated as the proportion of left choices made over the preceding 10 trials. (D) Correct rate around reversals during the last day of the DRL task and the three PRL tasks in WT mice. Data were analyzed using a mixed-effects model with trial, task, and their interaction as fixed effects. n = 27 mice for DRL, PRL 80–20, and PRL 70–30; n = 26 mice for PRL 60–40. Data are presented as mean ± SEM. Panels A and B were created with Biorender.

In the DRL and PRL tasks, the panel associated with the higher reward probability was reversed after 12 correct choices within the preceding 15 trials. Behavioral flexibility was assessed by measuring the correct rate, defined as the proportion of choices directed toward the panel with the higher reward probability, around each reversal (**Figure 1C**). Correct rate decreased immediately after reversal and gradually recovered as mice adapted to the new reward contingency (**Figure 1D**). This recovery became progressively slower as reward uncertainty increased, with the slowest adaptation observed in the PRL 60–40 task. These results demonstrate that the task sequence enables quantitative assessment of behavioral flexibility under varying levels of reward uncertainty.

### Premature responses are negatively associated with the initial acquisition of deterministic reversal learning

Premature responding is commonly used as a measure of impulsive behavior. We therefore quantified premature responses, defined as touches to the side panels before trial initiation by touching the initiation cue, during the DRL and PRL tasks (**Figure 2A**). Of note, premature responses were not counted as choices and were therefore excluded when calculating the correct rate. In wild-type (WT) mice, premature response count gradually decreased as the correct rate increased during DRL training (**Figure 2B, C**). Premature response count was significantly lower on the last day of the DRL task than on the first day (**Figure 2D**), indicating that mice progressively suppressed premature responses as they acquired the task.

**Figure 2.**
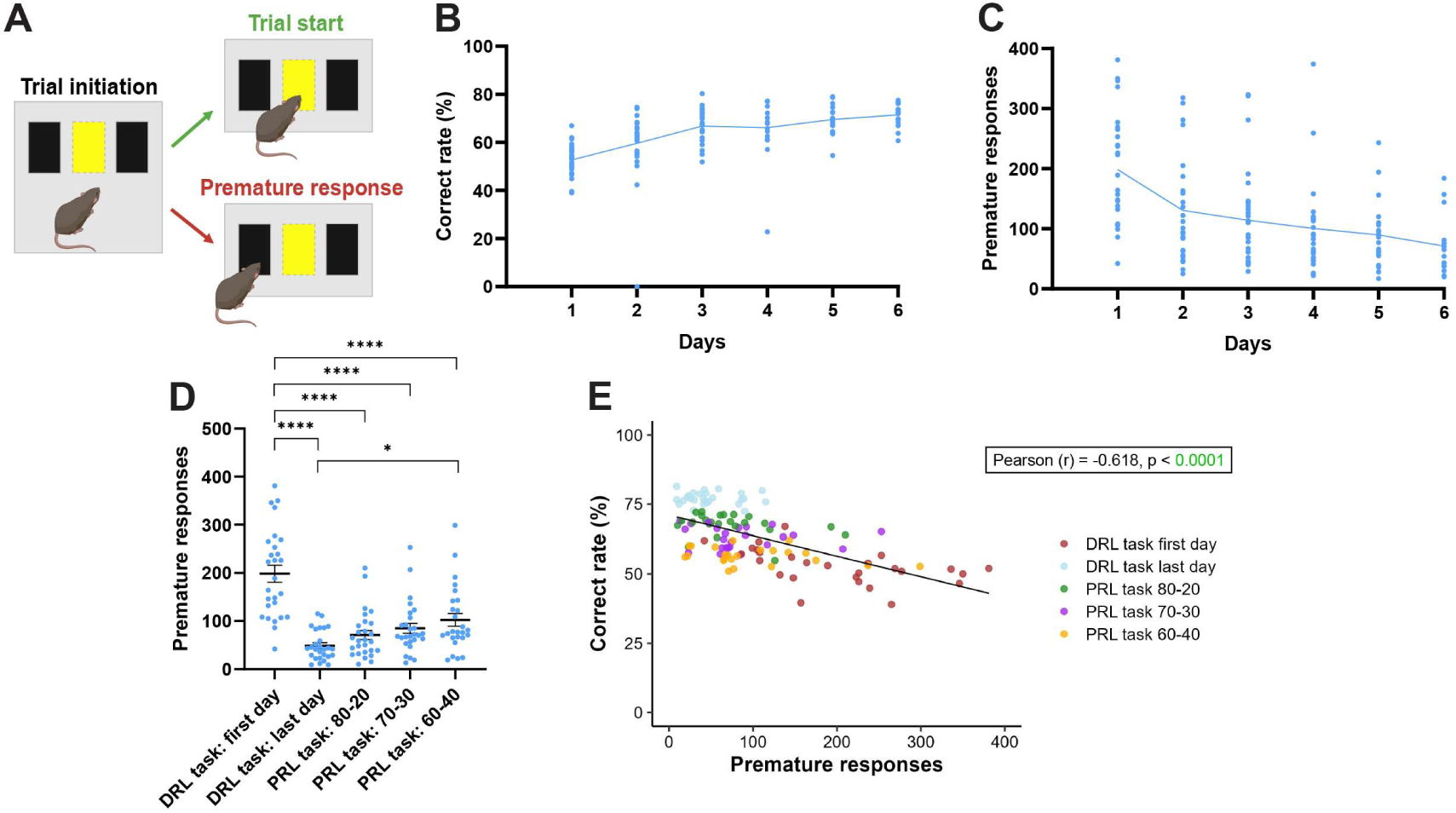
Relationship between premature responses and task performance in WT male mice. (A) Schematic illustrating premature responses during the DRL and PRL tasks. Each trial was initiated by a center-panel touch, followed by a choice of the left or right panel. Premature responses were defined as touches to the left or right panel before initiation of a trial by touching the center panel. (B) Premature response count during 6 days of DRL training. Data were analyzed using a linear mixed-effects model with day as a fixed effect. n = 27 mice. Points represent individual mice, and the line indicates the mean. (C) Correct rate during 6 days of DRL training. Data were analyzed using a linear mixed-effects model with day as a fixed effect. n = 27 mice. Points represent individual mice, and the line indicates the mean. (D) Premature response count on the first and last days of the DRL task and during the PRL 80–20, 70–30, and 60–40 tasks. Data were analyzed using a mixed-effects model with task as a fixed effect, followed by Tukey’s multiple-comparisons test. n = 27 mice for DRL, PRL 80–20, and PRL 70–30; n = 26 mice for PRL 60–40. (E) Relationship between premature response count and correct rate on the first and last days of the DRL task and during the PRL 80–20, 70–30, and 60–40 tasks. Each point represents one mouse, and colors indicate different task conditions. Pearson’s correlation coefficient (r) and p values were calculated separately for each condition.

We next examined how premature responses changed as reward uncertainty increased. Premature response count increased as reward contingencies became more uncertain, reaching its highest level during the PRL 60–40 task (**Figure 2D**). Thus, premature responses decreased with acquisition of deterministic reversal learning but increased under greater reward uncertainty.

We then asked whether premature responses were associated with task performance. Across the DRL and PRL tasks, premature response count was negatively associated with correct rate (**Figure 2E**). When we assessed the correlation for each task, this relationship was significant on the first day of the DRL task (**Figure 2E**). These findings indicate that greater impulsivity, represented as premature responses, is associated with poorer task performance, particularly during the initial acquisition of deterministic reversal learning.

### Optimization of reinforcement-learning models to assess adaptive decision-making

To characterize latent cognitive processes during reversal learning, we fitted behavioral data using five reinforcement-learning models described by Hattori et al^12^. These models were based on Q-learning and estimated trial-by-trial values of the chosen (Q_ch_) and unchosen (Q_unch_) options. Q_ch_ was updated using separate learning rates for rewarded and unrewarded outcomes (α_rew_ and α_unrew)_, whereas Q_unch_ was updated according to the forgetting rate of the unchosen option (δ). Choice probability was determined by the difference in action values, together with the sensitivity to this value difference (β_ΔQ)_ and a choice-bias parameter (β_o_). The five models differed in the combination of these parameters included (**Figure 3A**; **Table 1**), and we used a logistic-regression model for comparison.

**Figure 3.**
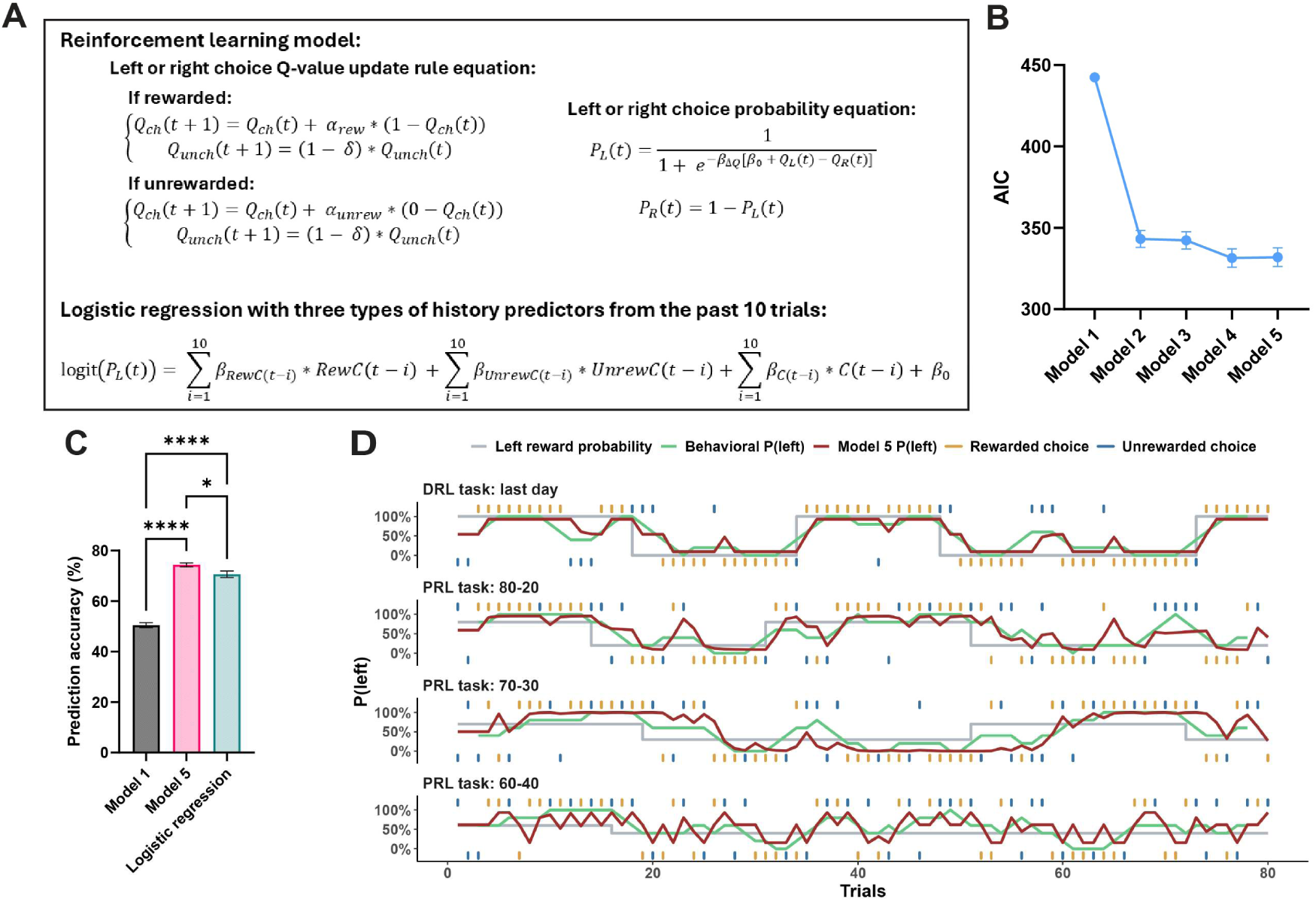
Optimization of reinforcement-learning models to assess adaptive decision-making. (A) Schematic of the five-parameter reinforcement-learning (RL) model and the 31-parameter logistic-history model. Q_ch_ and Q_unch_ denote the values of the chosen and unchosen options during value updating, respectively, whereas Q_L_ and Q_R_ denote the corresponding left and right action values used to calculate choice probability. (B) AIC values for five RL models fitted to WT datasets, which included first 80 trials from the last day of the DRL, PRL 80–20, and PRL 70–30 tasks. (C) Prediction accuracy of RL Model 1, RL Model 5, and a logistic-history model. Analyses used the first 80 trials from the last day of DRL and the PRL 80–20 and PRL 70–30 tasks. For the logistic-history model, the first 10 trials of each session were used to establish choice history and were excluded from prediction. Prediction accuracy was assessed by two-fold cross-validation. One-way ANOVA followed by Tukey’s multiple-comparisons test. (D) Observed and model-predicted probabilities of choosing the left panel across trials in a representative WT mouse during the last day of DRL, PRL 80–20, PRL 70–30, and PRL 60–40 tasks. The gray line indicates the reward probability assigned to the left panel. The green line indicates observed behavior, and the red line model-predicted behavior. Yellow and gray ticks indicate rewarded and unrewarded choices, respectively. n = 26 mice. Data are presented as mean ± SEM. * p < 0.05, ****p < 0.0001.

**Table 1.** Parameters included in the five reinforcement-learning models. Checkmarks indicate parameters included in each model. α_rew_ and α_unr_ denote learning rates following rewarded and unrewarded outcomes, β_ΔQ_ denotes sensitivity to the difference between action values, β₀ denotes constant choice bias and δ denotes forgetting of the unchosen option.

| Model | $\alpha_{\text{rew}}$ | $\alpha_{\text{unr}}$ | $\beta_{\Delta Q}$ | $\beta_0$ | $\delta$ | Interpretation |
| --- | --- | --- | --- | --- | --- | --- |
| 1 | — | — | — | ✓ | — | Constant side bias benchmark |
| 2 | ✓ | ✓ | ✓ | — | — | Outcome-specific value updating |
| 3 | ✓ | ✓ | ✓ | ✓ | — | Model 2 with side bias |
| 4 | ✓ | ✓ | ✓ | — | ✓ | Model 2 with forgetting |
| 5 | ✓ | ✓ | ✓ | ✓ | ✓ | Full model with bias and forgetting |

Model parameters were estimated from WT datasets from the last day of the DRL task and the PRL 80–20 and 70–30 tasks. We used the first 80 trials of each task, which all mice completed. The PRL 60–40 task was excluded because reversal performance was highly variable under this high-uncertainty condition, with some mice completing very few reversals. Among the reinforcement-learning models, Models 4 and 5 yielded similarly low Akaike Information Criterion (AIC) values (**Figure 3B**), indicating that inclusion of the additional side-bias parameter in Model 5 did not substantially improve model fit. We nevertheless used Model 5 for subsequent analyses because it represents the full model. Model 5 also showed higher cross-validated prediction accuracy than the logistic-history model (**Figure 3C**). The fitted model broadly reproduced the observed choice patterns, including in the PRL 60–40 task, which was not included in parameter estimation (**Figure 3D**).

### A subset of *Mecp2^+/−^* mice shows impaired acquisition of the deterministic reversal learning task with reduced engagement and increased premature responding

Having established a task sequence for adaptive decision-making, we examined *Mecp2*^+/−^ mice alongside their littermate WT controls. We used 8-week-old female mice, which exhibit only mild neurological deficits, to detect early signs of cognitive abnormalities^21^. Performance in the VD task was comparable between genotypes, with no differences in the number of days required to reach criterion or the proportion of mice progressing to the next task (**Figure 4–figure supplement 1**). In contrast, *Mecp2*^+/−^ mice were impaired in the DRL task. All WT mice reached the predefined criterion (correct rate >75% or >70% for 2 days) within 8 days, whereas only 52.4% of *Mecp2*^+/−^ mice reached the criterion within 14 days, resulting in a significantly lower progression rate to the PRL tasks (**Figure 4A**). We therefore classified *Mecp2^+/-^* mice as progressors and non-progressors and compared their results with those of WT mice.

**Figure 4.**
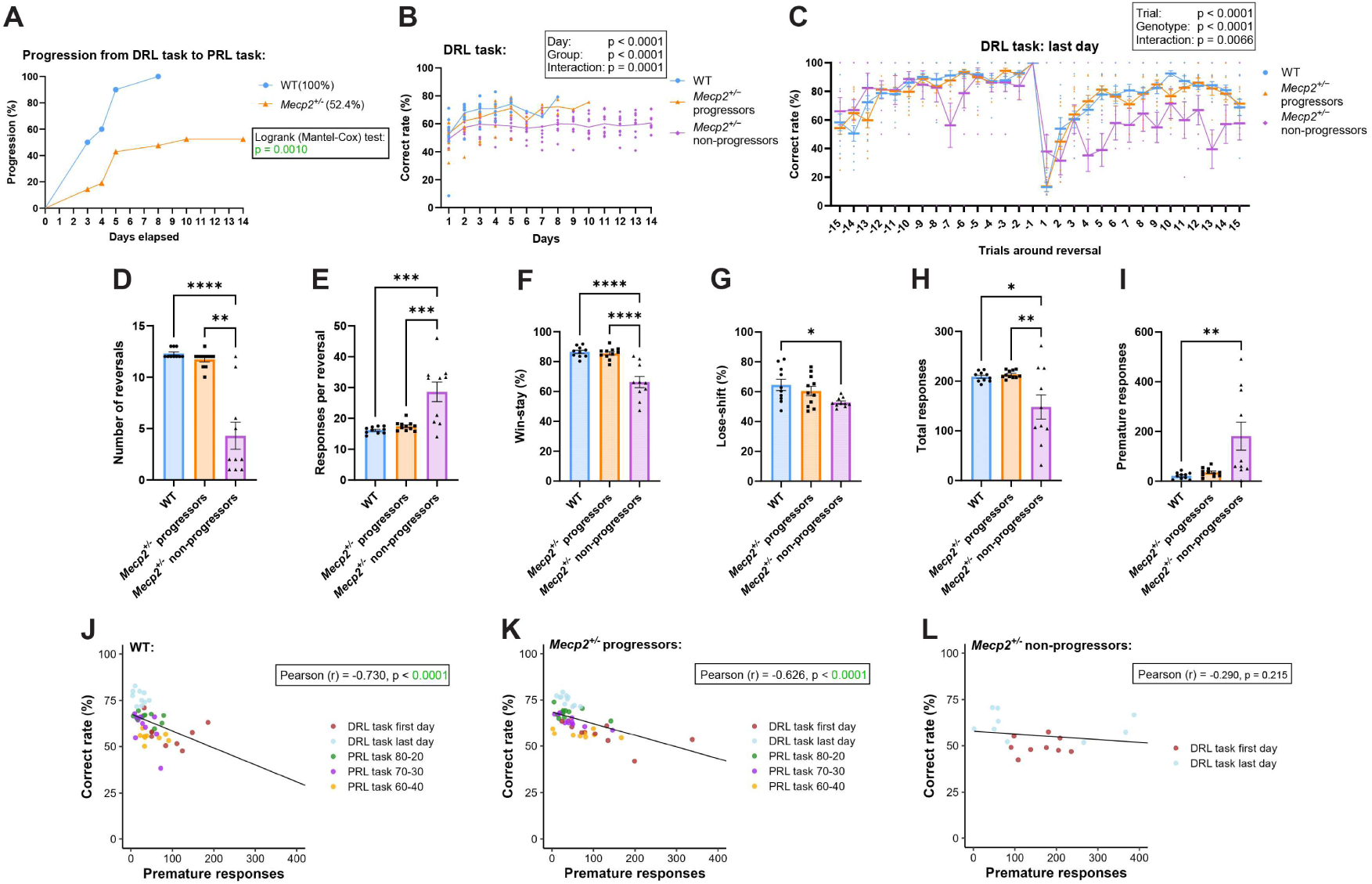
*Mecp2^+/−^* non-progressors exhibit impaired acquisition of the deterministic reversal learning task, reduced engagement, and increased premature responding. (A) Progression from the DRL task to the PRL tasks. Log-rank (Mantel–Cox) test. n = 10 WT and 21 *Mecp2*^+/−^ mice. (B) Correct rate during 14 days of DRL training. Data were analyzed using a linear mixed-effects model with day (categorical variable), group, and their interaction as fixed effects. Points represent individual mice, and the line indicates the mean. (C) Correct rate around reversals on the last day of the DRL task. For each mouse, correct rates at each trial relative to reversal were averaged during all reversals. Mixed-effects model with trial, group, and their interaction as fixed effects. (D) Number of reversals completed on the last day of the DRL task. Kruskal-Wallis test with Dunn’s multiple comparisons. (E) Responses per reversal on the last day of the DRL task. Data were analyzed using one-way ANOVA followed by Tukey’s multiple-comparisons test. (**F, G**) Win-stay (F) and lose-shift (G) probabilities on the last day of the DRL task. One-way ANOVA followed by Tukey’s multiple-comparisons test. (H) Total responses on the last day of the DRL task. One-way ANOVA followed by Tukey’s multiple-comparisons test. (I) Premature responses on the last day of the DRL task. Kruskal–Wallis test followed by Dunn’s multiple-comparisons test. (**J, K, L**) Relationship between premature responses and correct rate during the first and last days of the DRL task and PRL tasks in WT mice (J), *Mecp2*^+/−^ progressors (K) and *Mecp2*^+/−^ non-progressors (L). Each point represents one mouse, and colors indicate different task conditions. Pearson’s correlation coefficient (*r*) and *p* values were calculated separately for each condition. *n* = 10 WT mice, 11 *Mecp2*^+/−^ progressors, and 10 *Mecp2*^+/−^ non-progressors. Data are presented as mean ± SEM. *p < 0.05, **p < 0.01, ***p < 0.001, ****p < 0.0001.

As expected, *Mecp2*^+/−^ non-progressors exhibited impaired acquisition of the DRL task (**Figure 4B**). On the last day of the DRL task, *Mecp2*^+/−^ non-progressors showed impaired adaptation following reversals (**Figure 4C**), accompanied by fewer reversals and increased responses per reversal (**Figure 4D, E**). We further examined feedback-dependent choice strategies using win-stay and lose-shift probabilities. Win-stay probability is the likelihood of repeating the same choice after a rewarded outcome, whereas lose-shift probability is the likelihood of switching to the alternative choice after an unrewarded outcome. Both win-stay and lose-shift probabilities were reduced in non-progressors (**Figure 4F, G**), indicating altered feedback-dependent choice behavior. These alterations were accompanied by reduced task engagement, reflected by fewer total responses, and increased premature responses (**Figure 4H, I; Figure 4–figure supplement 2**).

Interestingly, the relationship between premature responses and correct rate differed significantly among groups (premature responses × group: F_2,111.4 =_ 25.66, p < 0.0001; **Figure 4J–L**). The negative association was strongest in WT mice (slope = −0.081), weaker in *Mecp2*^+/−^ progressors (slope = −0.043), and absent in *Mecp2*^+/−^ non-progressors (slope = −0.011). Pairwise comparisons confirmed significant differences in slopes between WT mice and progressors (p = 0.0015), WT mice and non-progressors (p < 0.0001), and progressors and non-progressors (p = 0.010). Thus, the association between premature responding and task performance became progressively weaker from WT mice to *Mecp2*^+/−^ progressors and non-progressors.

### *Mecp2*^+/−^ progressors exhibit normal probabilistic reversal learning with altered feedback strategy

In contrast to non-progressors, *Mecp2*^+/−^ progressors that successfully acquired the DRL task performed comparably to WT mice during both deterministic and probabilistic reversal learning (**Figures 4 and 5**). Correct rate around reversals, reversal number, and the responses per reversal were also comparable between genotypes across tasks (**Figure 5A–E**). However, *Mecp2*^+/−^ progressors exhibited an altered feedback-dependent choice strategy. In WT mice, win-stay probability decreased as reward uncertainty increased (Mixed-effects analysis: Tasks: p < 0.0001; **Figure 5F**). Win-stay probability similarly decreased with increasing uncertainty in *Mecp2*^+/−^ progressors but remained higher than in WT mice during the PRL 80–20 and 70–30 tasks (**Figure 5F**). In contrast, lose-shift probability was comparable between genotypes across tasks (**Figure 5G**). Thus, *Mecp2*^+/−^ progressors exhibited largely preserved behavioral flexibility under probabilistic reward conditions but showed an increased tendency to repeat choices following positive outcome.

**Figure 5.**
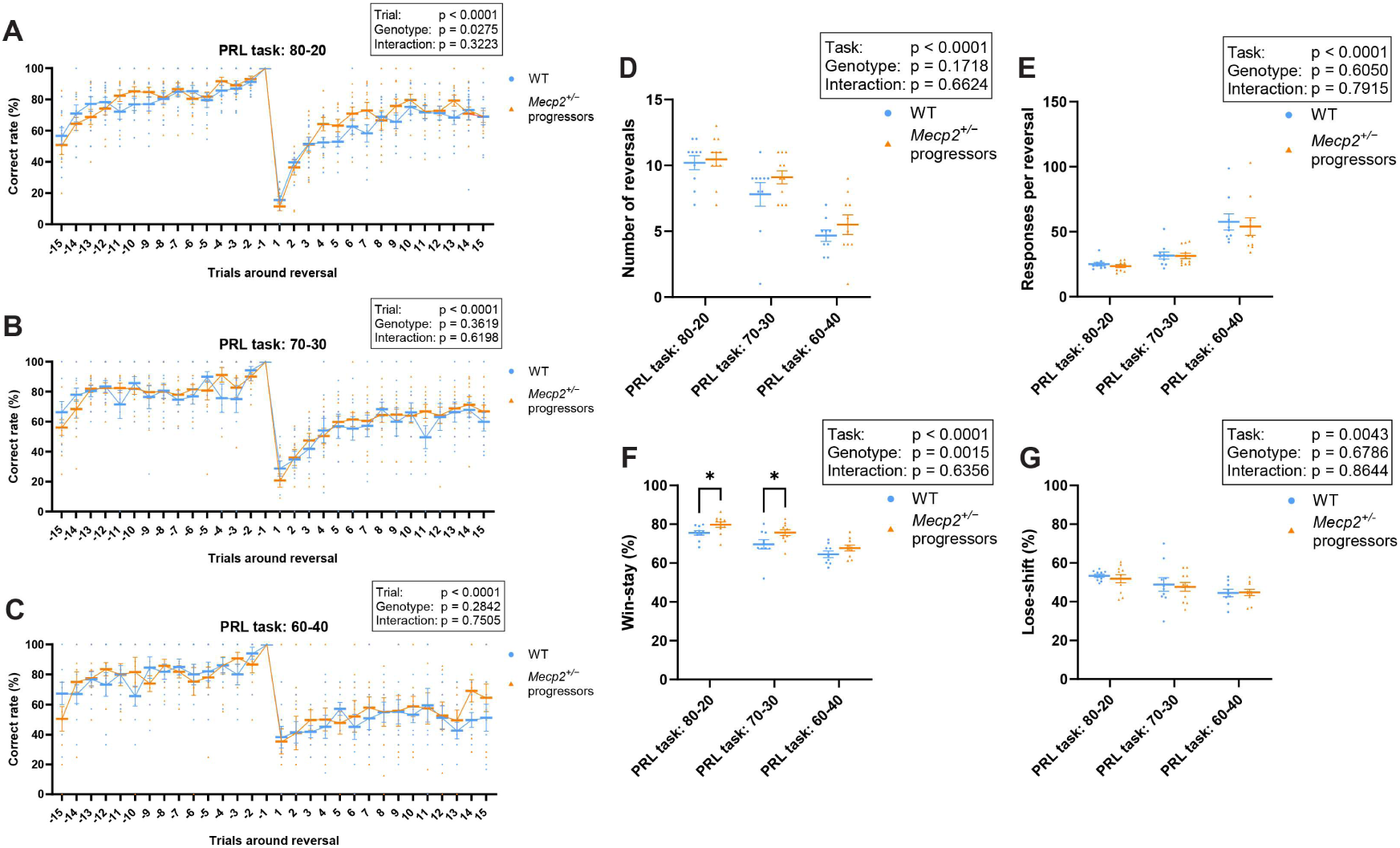
*Mecp2*^+/−^progressors exhibit normal probabilistic reversal learning with altered feedback strategy. (**A–C**) Correct rate around reversals during the PRL 80-20 (A), PRL 70-30 (B), and PRL 60-40 (C) tasks. Two-way repeated measures ANOVA. Fixed effects: trial, genotype, interaction. (**D–G**) Number of reversals (D), responses per reversal (E), win-stay probabilty (F), and lose-shift probability (G) during the PRL tasks. Mixed-effects analysis with Tukey’s multiple comparisons test. Fixed effects: task, genotype, interaction. For WT mice, *n* = 10 for the first and last days of DRL, PRL 80–20, and PRL 70–30, and *n* = 9 for PRL 60–40. For *Mecp2*^+/−^ progressors, *n* = 11 for the first and last days of DRL, PRL 80–20, and PRL 70–30, and *n* = 10 for PRL 60–40. Points represent individual mice, and the line and error bars indicate mean ± SEM. \**p* < 0.05.

### *MECP2-*Tg1 mice exhibit mild impairment during deterministic reversal learning but normal probabilistic reversal learning

Overexpression of MeCP2 is associated with *MECP2* duplication syndrome, whose neurological features are recapitulated in *MECP2*-Tg1 mice^19^. To examine the effect of MeCP2 overexpression on adaptive decision-making, we tested 8-week-old male *MECP2-*Tg1 mice together with their littermate WT controls. Males were used because *MECP2* duplication syndrome primarily affects males, and the age was matched to that of the *Mecp2*^+/−^ cohort.

Performance in the VD task was comparable between genotypes, with all mice reaching criterion within 5 days (**Figure 6–figure supplement 1A–C**). During DRL training, *MECP2*-Tg1 mice exhibited a mild but significant reduction in correct rate compared with WT mice, despite progressive improvement across training days (**Figure 6A**). However, the number of days required to reach criterion and the proportion of mice progressing to the PRL tasks did not differ between genotypes, indicating that overall acquisition of deterministic reversal learning was largely preserved (**Figure 6–figure supplement 1D, E**).

**Figure 6.**
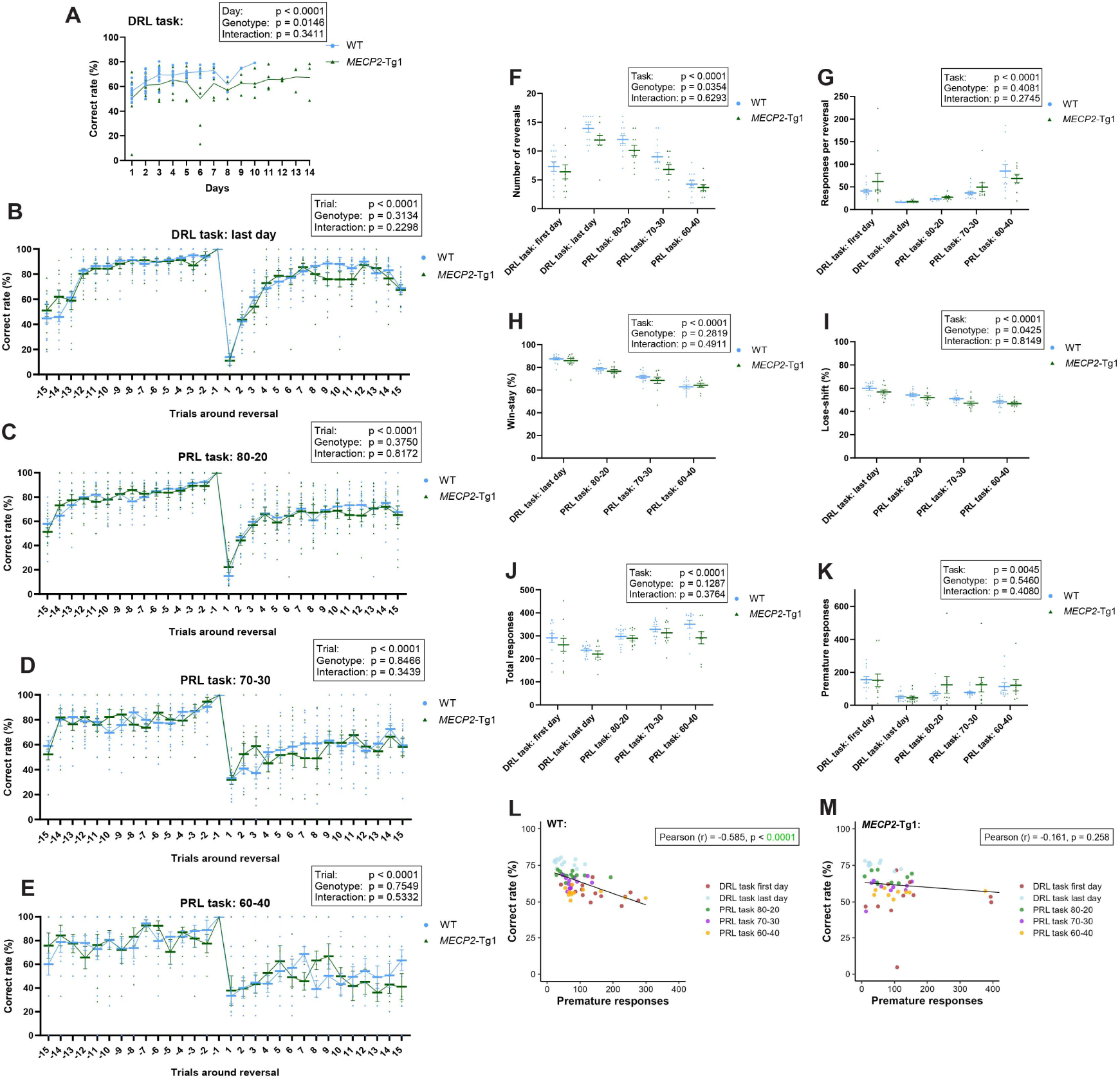
*MECP2-*Tg1 mice exhibit mild impairment during deterministic reversal learning but normal probabilistic reversal learning. (A) Correct rate during 14 days of DRL training. Data were analyzed using a linear mixed-effects model with day (categorical variable), group, and their interaction as fixed effects. (B) Correct rate around reversals on the last day of the DRL task. Mixed-effects analysis with trial, genotype, and their interaction as fixed effects. (**C–E**) Correct rate around reversals during the PRL 80–20 (C), PRL 70–30 (D), and PRL 60–40 (E) tasks. Two-way repeated-measures ANOVA with trial, genotype, and their interaction as fixed effects. (**F–K**) Number of reversals (F), responses per reversal (G), win-stay probability (H), lose-shift probability (I), total responses (J), and premature responses (K) during the DRL and PRL tasks. Mixed-effects analysis with task, genotype, and their interaction as fixed effects, followed by Tukey’s multiple-comparisons test. (**L, M**) Relationship between premature responses and correct rate during the first and last days of the DRL task and the PRL 80–20, 70–30, and 60–40 tasks in WT (L) and *MECP2*-Tg1 (M) mice. Each point represents one mouse, and colors indicate different task conditions. Pearson’s correlation coefficient (*r*) and *p* values were calculated separately for each condition. For WT mice, *n* = 13 for the first and last days of DRL, PRL 80–20, and PRL 70–30, and *n* = 12 for PRL 60–40. For *MECP2*-Tg1 mice, *n* = 11 for the first and last days of DRL, *n* = 10 for PRL 80–20 and PRL 70–30, and *n* = 9 for PRL 60–40. Points represent individual mice, and lines and error bars indicate mean ± SEM.

Behavioral flexibility during the DRL and PRL tasks was also comparable between genotypes, as assessed by adaptation around reversals (**Figure 6B–E**), reversal number (**Figure 6F**), and responses per reversal (**Figure 6G**). Win-stay and lose-shift probabilities, task engagement, premature responding were also similar between *MECP2*-Tg1 and WT mice (**Figure 6H–K**; **Figure 6–figure supplement 1F, G**). Although the relationship between premature responses and correct rate appeared different between WT and *MECP2*-Tg1 mice (**Figure 6L, M**), a linear mixed-effects model revealed neither a significant overall association between premature responses and correct rate (F_1,104.2 =_ 1.78, p = 0.185) nor a significant interaction between premature responses and genotype (F_1,107.0 =_ 0.73, p = 0.393). Together, these findings indicate that two-fold MeCP2 overexpression produces a modest impairment during the acquisition of deterministic reversal learning but has little effect on subsequent behavioral flexibility, feedback-dependent choice strategies, task engagement, or premature responding at this age.

### *Kmt2c*^+/fs^ mice show largely intact deterministic and probabilistic reversal learning

We next examined *Kmt2c*^+/fs^ mice, which exhibit autism-related behavioral phenotypes^20^. Like *Mecp2*^+/−^ and *MECP2*-Tg1 mice, *Kmt2c*^+/fs^ mice did not differ from WT mice in acquisition or performance in the VD task (**Figure 7–figure supplement 1A–C**). Similarly, performance during DRL training was comparable between genotypes, with no differences in the number of days required to reach criterion, progression to the PRL tasks, or correct rate (**Figure 7A**; **Figure 7–figure supplement 1D, E**). Behavioral flexibility parameters, as assessed by correct rate around reversals, reversal number, responses per reversal, and win-stay and lose-shift probabilities, were also largely comparable across the DRL and PRL tasks (**Figure 7B–I**). Although *Kmt2c*^+/fs^ mice tended to require more responses per reversal during the PRL 60-40 task, this difference did not reach statistical significance (**Figure 7G**).

**Figure 7.**
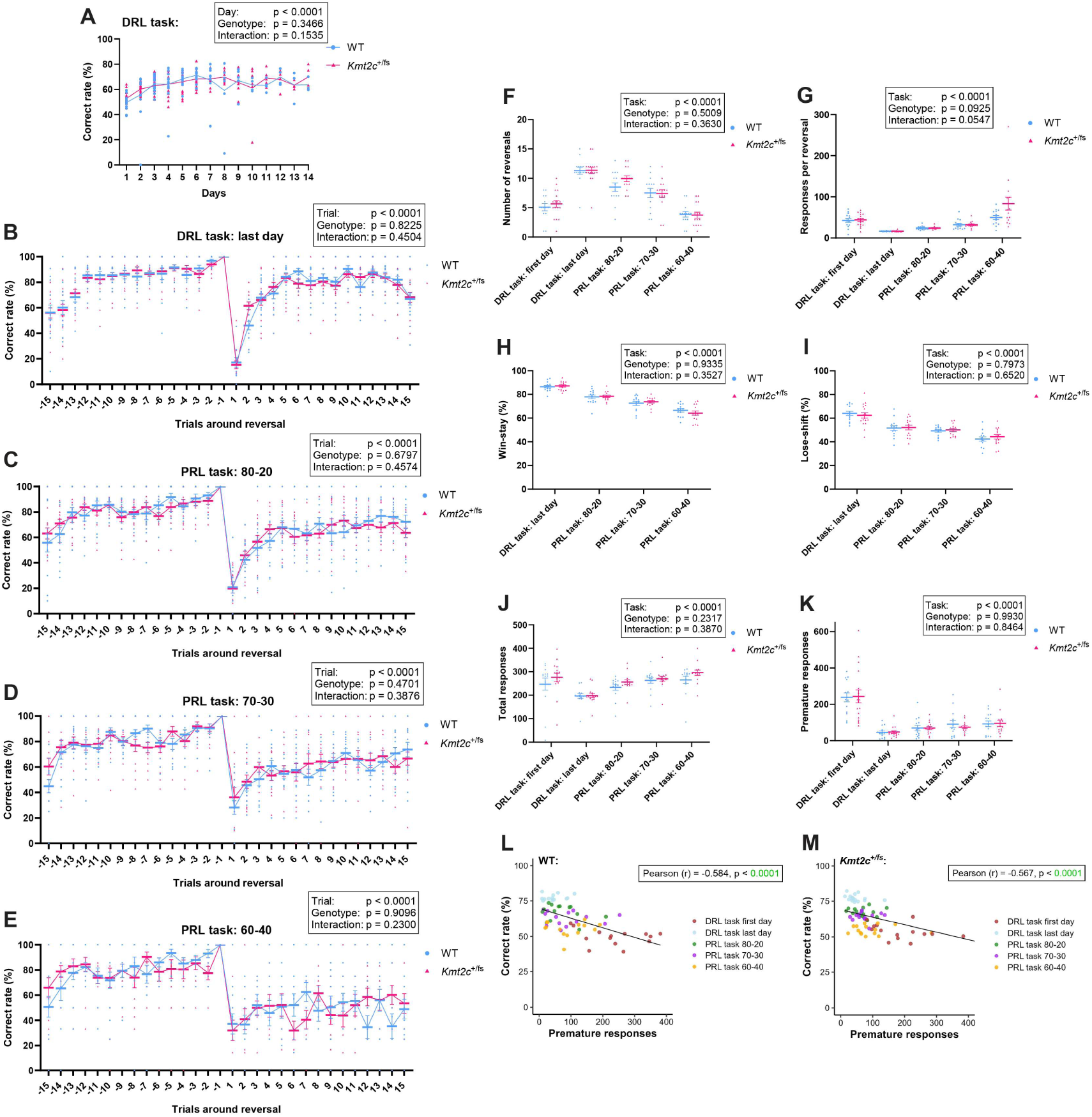
*Kmt2c*^+/fs^ mice exhibit largely normal deterministic and probabilistic reversal learning. (A) Correct rate during 14 days of DRL training. Data were analyzed using a linear mixed-effects model with day (categorical variable), group, and their interaction as fixed effects. (B) Correct rate around reversals on the last day of the DRL task. Mixed-effects analysis with trial, genotype, and their interaction as fixed effects. (**C–E**) Correct rate around reversals during the PRL 80–20 (C), PRL 70–30 (D), and PRL 60–40 (E) tasks. Two-way repeated-measures ANOVA with trial, genotype, and their interaction as fixed effects. (**F–K**) Number of reversals (F), responses per reversal (G), win-stay probability (H), lose-shift probability (I), total responses (J), and premature responses (K) during the DRL and PRL tasks. Mixed-effects analysis with task, genotype, and their interaction as fixed effects, followed by Tukey’s multiple-comparisons test. (**L, M**) Relationship between premature responses and correct rate during the first and last days of the DRL task and the PRL 80–20, 70–30, and 60–40 tasks in WT (L) and *Kmt2c^+/fs^* (M) mice. Each point represents one mouse, and colors indicate different task conditions. Pearson’s correlation coefficient (*r*) and *p* values were calculated separately for each condition. n = 14 WT mice for all tasks. For *Kmt2c^+/fs^* mice, n = 17 for the VD and DRL tasks, and n = 15 for the PRL tasks. Points represent individual mice, and lines and error bars indicate mean ± SEM. \**p* < 0.05.

Task engagement and premature responding were also comparable between genotypes across tasks (**Figure 7J, K; Figure 7–figure supplement 1F, G**). Across tasks, a linear mixed-effects model revealed a significant negative association between premature responses and correct rate (F_1,137.9 =_ 8.56, p < 0.01) and this relationship did not differ significantly between genotypes (premature responses x correct rate: F_1,135.8 =_ 1.21, p = 0.274; **Figure 7L, M**). Together, these findings indicate that *Kmt2c*^+/fs^ mice show largely preserved acquisition and behavioral flexibility across deterministic and probabilistic reversal learning.

### Reinforcement-learning modeling identifies altered value updating in *Kmt2c^+/fs^* mice

Across the three NDD models, conventional behavioral measures revealed distinct degrees of impairment. Deterministic reversal learning was significantly affected in a subset of *Mecp2*^+/−^ mice, whereas *MECP2*-Tg1 mice showed a mild impairment in the acquisition of the DRL task. Overall behavioral performance was largely preserved in *Kmt2c*^+/fs^ mice, although they showed a trend toward increased responses per reversal in the PRL 60–40 task.

To characterize decision-making processes not captured by these conventional measures, we applied the optimized reinforcement-learning model to datasets from *Mecp2^+/-^*, *MECP2*-Tg1, and *Kmt2c^+/fs^* mice. Because only half of the *Mecp2*^+/−^ mice progressed to the PRL tasks, the analysis was restricted to *Mecp2*^+/−^ progressors. Model parameters and AIC were compared with those of the corresponding littermate WT mice.

AIC was comparable between each mutant group and its corresponding WT controls, indicating similar model fits across genotypes (**Figure 8A–C**). Model parameters did not differ between *Mecp2*^+/−^ progressors and WT mice (**Figure 8A**), consistent with their largely preserved performance in the DRL and PRL tasks. Similarly, *MECP2*-Tg1 mice showed parameters comparable to those of WT mice (**Figure 8B**).

**Figure 8.**
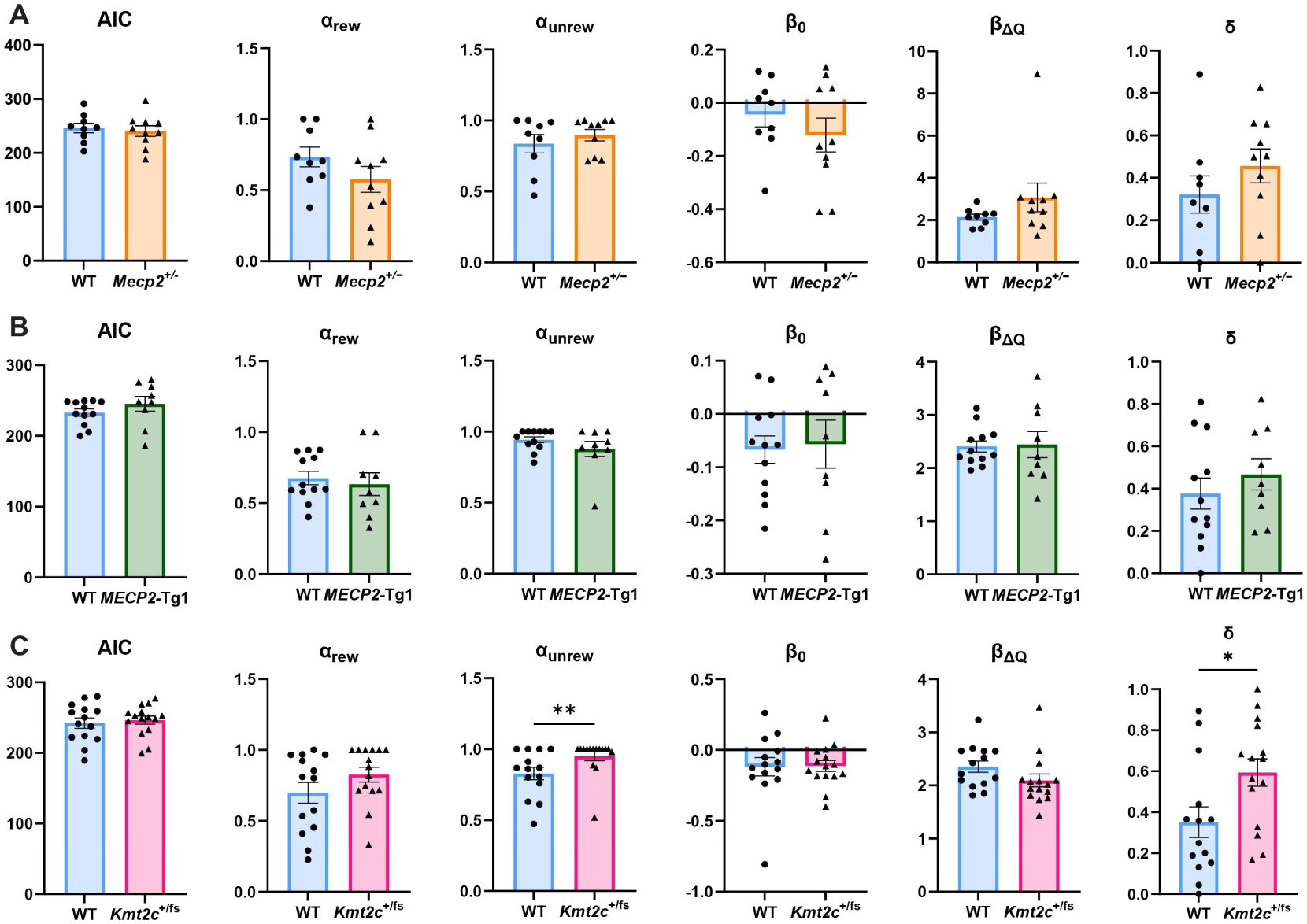
Reinforcement-learning modeling reveals altered value updating and forgetting in *Kmt2c*^+/fs^ mice. (A) AIC values and five parameters (α_rew_, α_unrew_, β_Q_, β_0_, and δ) obtained from reinforcement-learning models in WT and *Mecp^2^*^+/−^ mice. n = 9 WT and 10 *Mecp2*^+/−^ mice. (B) AIC values and five parameters obtained from reinforcement-learning models in WT and *MECP2*-Tg1 mice. n = 12 WT and 9 *MECP2*-Tg1 mice. (C) AIC values and five parameters obtained from reinforcement-learning models in WT and *Kmt2c^+/fs^* mice. *n* = 14 WT and 15 *Kmt2c^+/fs^* mice. AIC was compared between genotypes using two-way repeated-measures ANOVA followed by Tukey’s multiple-comparisons test. Parameters were analyzed using Mann–Whitney tests. Bars and error bars indicate mean ± SEM. * p < 0.05, ** p < 0.01, **** p < 0.0001.

In contrast, *Kmt2c^+/fs^* mice showed selective alterations in model parameters. The learning rate following unrewarded outcomes (α_unrew)_ was significantly higher in *Kmt2c^+/fs^* mice than in WT mice, whereas the learning rate following rewarded outcomes (α_rew)_ was unchanged (**Figure 8C**). *Kmt2c^+/fs^* mice also exhibited an increased forgetting rate for unchosen options (δ). Thus, *Kmt2c^+/fs^* mice updated the values of chosen options more strongly following unrewarded outcomes and more rapidly discounted the values of unchosen options.

To determine how changes in individual model parameters influence behavioral performance, we conducted simulations in which each of the five parameters was systematically varied while the remaining parameters were fixed at the estimates obtained for individual WT mice. For each parameter setting, behavioral performance was simulated across the DRL and PRL tasks, and correct rate, reversal number, and responses per reversal were quantified. The simulations revealed distinct effects of the model parameters on behavioral performance (**Figure 9A–E**). Increasing α_rew_, which determines the extent to which action values are updated following rewarded outcomes, generally improved performance, resulting in higher correct rates and reversal numbers and fewer responses per reversal across tasks (**Figure 9A**). Increasing β_ΔQ_, which determines the sensitivity of choices to differences in estimated action values, similarly improved performance (**Figure 9D**). β_0_, representing side bias, also affected simulated performance; however, this effect likely reflects the task design, as the left panel was designated as the initially correct option for all tasks (**Figure 9E**).

**Figure 9.**
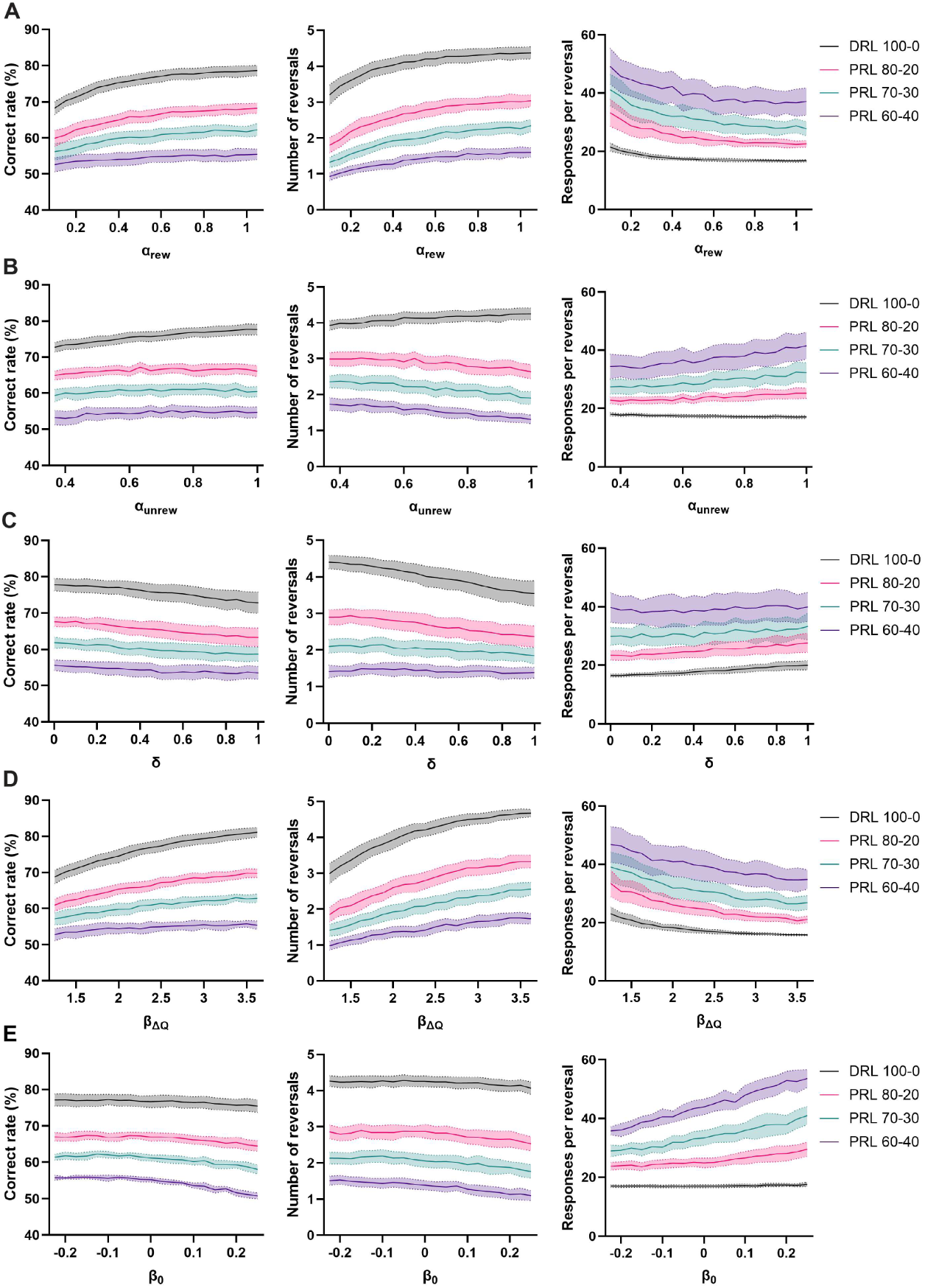
Reinforcement-learning parameters differentially affect simulated task performance. (**A–E**) Effects of α_rew (_**A**), α_unrew (_**B**), δ (**C**), β_ΔQ (_**D**), and β_0 (_**E**) on correct rate, number of reversals, and responses per reversal across the DRL and PRL tasks. For each parameter value, 100 simulations were performed for each of 14 WT mice, with the remaining parameters fixed at the estimates for each mouse. Lines indicate means across mice after averaging the 100 simulations within each mouse; shaded areas indicate 95% confidence intervals. Data were analyzed using linear mixed-effects models with task, the indicated model parameter, and their interaction as fixed effects and mouse identity as a random effect. For all three performance measures, significant effects of task, parameter, and task × parameter interaction were observed for α_rew_, α_unrew_, β_ΔQ_, and β_0 (_all *p* < 0.0001). For δ, effects of task and parameter were significant for all performance measures (all *p* < 0.0001), as were task × δ interactions (*p* < 0.0001 for correct rate and number of reversals; *p* < 0.01 for responses per reversal).

Notably, α_unrew_ and δ, both of which were increased in *Kmt2c*^+/fs^ mice, exerted distinct and partially opposing effects on task performance, and these effects varied with reward uncertainty. In the DRL task, increasing α_unrew_ increased correct rate and reversal number (**Figure 9B**), whereas increasing δ reduced these measures (**Figure 9C**). The beneficial effect of increased α_unrew_ diminished as reward uncertainty increased, with higher α_unrew_ instead associated with reduced reversal efficiency, reflected by increased responses per reversal in the PRL tasks (**Figure 9B**). Thus, the simultaneous increase in α_unrew_ and δ may partially offset their effects on the overall performance under deterministic conditions, whereas such compensation may diminish as reward uncertainty increases.

Together, these simulations indicate that individual reinforcement-learning parameters have dissociable effects on task performance and suggest that apparently preserved behavioral performance in *Kmt2c^+^*^/fs^ mice may arise in part from the opposing effects of increased updating following unrewarded outcomes and increased forgetting of unchosen options. In addition, the simulations showed that the effect of increased α_unrew_ became less beneficial as reward uncertainty increased, with higher α_unrew_ associated with more responses per reversal in the probabilistic tasks. This finding is consistent with the trend toward increased responses per reversal observed in *Kmt2c*^+/fs^ mice in the PRL 60–40 task.

## Discussion

In this study, we developed a sequence of operant conditioning tasks to assess multiple components of adaptive decision-making and combined it with reinforcement-learning models to infer latent cognitive processes. Application of this framework to three mouse models revealed distinct behavioral phenotypes and identified altered value updating in *Kmt2c^+/fs^* mice that was not readily apparent from task performance. These findings demonstrate the utility of our framework for characterizing distinct cognitive phenotypes in mouse models of NDDs.

*MECP2* encodes methyl-CpG-binding protein 2 (MeCP2), a chromatin-associated protein that regulates chromatin organization and neuronal gene expression^22,23^. Loss of MeCP2 function causes Rett syndrome^24,25^, a neurodevelopmental disorder that primarily affects females, and *Mecp2*^+/−^ female mice reproduce its core neurological symptoms^18^. Although these mice have been extensively studied, their relatively mild and variable phenotype at young ages can limit the detection of early cognitive abnormalities^21^. Here, we detected a significant impairment in the acquisition of deterministic reversal learning in a subset of *Mecp2*^+/−^ females at 8 weeks of age, when overt neurological symptoms remain relatively mild. The severity of this impairment was highly variable, separating the mice into progressors and non-progressors. Such variability may in part reflect differences in X-chromosome inactivation and the resulting mosaic expression of MeCP2 across neural circuits^26^. Interestingly, *Mecp2*^+/−^ progressors that successfully acquired deterministic reversal learning performed largely comparably to WT mice under probabilistic reward conditions, despite an increased win-stay probability. By contrast, impaired acquisition of the DRL task in non-progressors was accompanied by decreased win-stay and lose-shift probabilities, increased premature responding, and reduced task engagement. Although these findings are consistent with impaired inhibitory control and task engagement, future studies incorporating independent measures of attention, motivation, and impulse control will be necessary to determine the relative contribution of these processes.

*MECP2* gene is dosage-sensitive, and increased *MECP2* dosage causes *MECP2* duplication syndrome, a severe neurodevelopmental disorder primarily affecting males that is modeled by *MECP2*-Tg1 mice^19,27^. In contrast to *Mecp2*^+/−^ female mice, MeCP2 overexpression in *MECP2*-Tg1 mice produced only mild alterations at 8 weeks of age. Previous studies have shown that many phenotypes in *MECP2*-Tg1 mice become more pronounced after 10 weeks of age^19,28^, raising the possibility that the relatively preserved cognitive performance observed here reflects the young age at testing. It will therefore be important to determine whether deficits in adaptive decision-making emerge at later ages as disease phenotypes progress.

Loss-of-function variants in *KMT2C* (Lysine methyl transferase 2c), which encodes a histone methyltransferase involved in transcriptional regulation through H3K4 methylation, have been associated with neurodevelopmental disorder, including autism spectrum disorders and Kleefestra syndrome 2^29,30^. A previous study demonstrated that *Kmt2c*^+/fs^ mice exhibit several autism-related phenotypes, including reduced sociality and impaired working memory^20^. The same study also found reduced flexibility in the Barnes maze and IntelliCage tests^20^, in which *Kmt2c*^+/fs^ mice showed increased time spent around previously correct choices. In contrast, in the present study, *Kmt2c*^+/fs^ mice showed largely preserved behavioral flexibility in the DRL and PRL tasks. In the IntelliCage test, the correct location was changed daily, whereas reversals in the present DRL and PRL tasks typically occurred within minutes. The Barnes maze phenotype was detected in a probe test following spatial learning over multiple days, rather than during repeated within-session reversals. Thus, these paradigms may assess behavioral adaptation over substantially different timescales.

Importantly, computational modeling uncovered alterations in latent value-updating processes in *Kmt2c*^+/fs^ mice despite relatively preserved performance in the reversal learning tasks, illustrating how computational approaches can reveal cognitive abnormalities that are not readily captured by conventional measures. Specifically, *Kmt2c*^+/fs^ mice showed increases in α_unrew_ and δ, consistent with stronger updating following unrewarded outcomes and faster forgetting of unchosen options. To better understand the functional consequences of these parameter changes, we performed simulations and found that learning rates for rewarded and unrewarded outcomes had distinct effects on behavioral performance. Increasing α_rew_ generally improved performance, whereas increasing α_unrew_ had opposite effects under deterministic and probabilistic reward contingencies. Under deterministic conditions, stronger updating following an unrewarded outcome facilitated adaptation because the absence of reward reliably indicated an incorrect choice. In contrast, under probabilistic conditions, an unrewarded outcome could represent misleading feedback rather than a change in the underlying reward contingency, such that stronger updating could impair adaptive behavior. Previous computational studies in individuals with autism have also suggested that learning from rewarded and unrewarded outcomes can differentially contribute to adaptive behavior, although the specific alterations differed from those observed here^7^.

Our simulations further showed that α_unrew_ and δ were associated with opposing effects on the performance of the DRL task. Increasing α_unrew_ improved task performance, whereas increasing δ impaired performance. This suggests that retaining the value of unchosen option benefits decision-making when reward contingencies are deterministic. The opposing effects of these parameters also raise the possibility that their simultaneous increase in *Kmt2c*^+/fs^ mice partially offset one another, contributing to the relatively preserved performance in the DRL task.

Taken together, our task sequence revealed distinct patterns of adaptive decision-making in the three NDD models rather than a common impairment in behavioral flexibility. In contrast to *Kmt2c*^+/fs^ mice, reinforcement-learning parameters were not significantly altered in either *Mecp2*^+/−^ or *MECP2*-Tg1 mice, suggesting that the computational alterations observed in *Kmt2c*^+/fs^ mice may arise from biological mechanisms distinct from those with altered MeCP2 dosage. In rodents, distinct orbitofrontal circuits contribute to learning from positive and negative outcomes during probabilistic reversal learning^31^, whereas the retrosplenial cortex persistently represents reward history and value-related information^12^. Although the neural processes underlying reinforcement-learning parameters remain unclear, combining behavioral modeling with neural recording and circuit manipulation may help identify the circuit mechanisms underlying cognitive alterations. The computational alterations identified in *Kmt2c*^+/fs^ mice may therefore provide well-defined phenotypes for investigating how molecular and cellular alterations affect neural circuits supporting adaptive decision-making.

There are several limitations in this study. First, the sequential nature of the task battery limited some comparisons: only mice that acquired DRL progressed to PRL, and this limitation was particularly relevant to *Mecp2*^+/−^ mice, approximately half of which did not progress to PRL. Second, PRL conditions were presented in a fixed order of increasing reward uncertainty, such that effects of uncertainty cannot be fully dissociated from task order and accumulated experience. Third, the present study focused on relatively young mice, which may limit detection of phenotypes that emerge with age. Longitudinal application of the same framework will be important for determining how decision-making phenotypes evolve during disease progression, particularly in *MECP2*-Tg1 mice. Finally, although our framework identified distinct behavioral and computational alterations across three NDD models, the mechanisms underlying these alterations remain unknown and require further investigation.

In conclusion, our findings demonstrate that combining quantitative behavioral assessment with computational modeling can distinguish specific components of cognitive dysfunction across genetically distinct mouse models. An important advantage of touchscreen cognitive testing is its potential for cross-species translation^32^. Extending this framework to additional cognitive domains, such as working memory^17^, may further enable multidimensional characterization of cognitive dysfunction and facilitate future cross-species studies.

## Materials and Methods

### Animals

Mouse experiments were conducted in accordance with the guidelines of the RIKEN Animal Experiment Committee. Mice were weaned at postnatal day 28 and housed with 3–5 littermates per cage. All mice were maintained on a C57BL/6 background. The following mouse lines were used in this study: Mecp2–heterozygous (*Mecp2^+/-^*)^18^, *MECP2*-Tg1^19^, and *Kmt2c^+/fs^* ^20^. *Mecp2^+/-^* females, *MECP2*-Tg1 males, and *Kmt2c^+/fs^* males were compared with age- and sex-matched wild-type mice. All mice were at least 2 months old at the time of the experiments. *Kmt2c^+/fs^* mice were either 8–9 or 11–15 weeks old at the start of experiments. The results from two age groups were combined because no statistically significant differences were detected.

### Operant House

All tasks were conducted during the night in the Operant House, a custom-designed home-cage system for automated operant conditioning (**Figure 1A**)^17^. Briefly, mice were individually housed in cages containing bedding with ad libitum access to food. Water access was restricted during the night and provided through the tasks. One wall of the cage was equipped with three panel windows and an LCD monitor that presented visual cues and detected touches through the panel windows. The opposite wall contained a water-delivery system consisting of a water slit and a retractable water bottle. A green LED signaled successful trial completion and access to the water bottle. Ceiling-mounted lights provided a light stimulus following an incorrect choice. Mice were monitored using a motion-detection system consisting of an infrared (IR) camera and IR lights. A Raspberry Pi 4 single-board computer controlled the system components and recorded behavioral events throughout the tasks.

### Task sequence for adaptive decision-making

The task sequence consisted of a 1-day habituation period followed by four task phases: pre-visual discrimination (Pre-VD), visual discrimination (VD), deterministic reversal learning (DRL), and probabilistic reversal learning (PRL) (**Figure 1B**). Each task was conducted either for a predetermined number of days or until mice reached a predefined performance criterion. Daily sessions ended when mice completed the required number of correct trials or 10 hours after the start of the session. During habituation, mice had free access to water during the 10-h nighttime session. During subsequent tasks, daily water access through task performance was limited to 160 seconds for mice weighing 20–25 g and 200 seconds for mice weighing 25–30 g. These durations were predetermined in pilot experiments to provide sufficient water intake to maintain body weight. If mice did not obtain the predetermined amount of water through tasks, supplemental water was provided on the following day.

#### Habituation

Mice were habituated in the Operant House for 1 day.

#### Pre-visual discrimination (Pre-VD)

In each trial, one of the three panels was randomly illuminated. Touching the illuminated panel resulted in 2 seconds of water access. The inter-trial interval (ITI) was 5 seconds. If a mouse did not touch the illuminated panel within 360 seconds, 1 second of water access was provided. The Pre-VD task was conducted for 1 day.

#### Visual discrimination (VD)

The VD task was performed as described for Pre-VD, except that touching an unilluminated panel resulted in flashing of the roof lights for 10 seconds. Water was available only through task performance. Mice continued the VD task until they achieved a correct rate ≥65% for two consecutive days.

#### Deterministic reversal learning (DRL)

Mice initiated each trial by touching the flashing center panel. Following trial initiation, the left and right panels were illuminated, with one panel designated as the correct choice. The left panel was initially designated as the correct choice. A correct choice resulted in 1 second of water access, whereas an incorrect choice resulted in flashing of the roof lights for 4 seconds. Thus, the reward contingencies were deterministic (100–0). The ITI was 3 seconds. The correct and incorrect sides were reversed after 12 correct choices within 15 consecutive trials. Mice continued DRL training until they achieved either a correct rate ≥70% for two consecutive days or ≥75% on a single day.

#### Probabilistic reversal learning (PRL)

Following acquisition of the DRL task, mice sequentially performed three PRL tasks with increasing reward uncertainty: PRL 80–20, PRL 70–30, and PRL 60–40. In PRL 80–20, choices of the correct panel resulted in water access with 80% probability and roof-light stimulation with 20% probability, whereas choices of the incorrect panel resulted in roof-light stimulation with 80% probability and water access with 20% probability. The corresponding probabilities were 70% and 30% in PRL 70–30 and 60% and 40% in PRL 60–40. The left panel was initially designated as the correct choice, and the correct side was switched to the other side after 12 correct choices within 15 consecutive trials. The ITI was 3 seconds, and each PRL task was conducted for 1 day.

### Task performance parameters

Task acquisition was assessed by the number of days required to reach the predefined criterion for the VD and DRL tasks and by the proportion of mice that reached each criterion within 14 days. Days to reach the VD criterion were defined as the number of days from the first day of VD training until mice achieved a correct rate ≥65% for two consecutive days. Days to reach the DRL criterion were defined as the number of days from the first day of DRL training until mice achieved either a correct rate ≥70% for two consecutive days or ≥75% on a single day. The proportion of mice progressing from VD to DRL and from DRL to PRL was calculated as:

*Progression (%) = (Number of mice that achieved the task within 14 days / Number of mice that started the task) * 100*

For subsequent behavioral analyses, the first and last days of DRL training and each of the three PRL tasks were analyzed. The first day of DRL represented initial acquisition of the reversal rule, whereas the last day represented performance after mice had acquired the task.

The following behavioral measures were quantified:

1. *Total responses* were defined as the total number of correct and incorrect panel touches during a session. Premature responses were excluded.
2. *Correct rate* was calculated as the percentage of responses directed toward the currently correct panel. Premature responses were excluded from this calculation.
3. *Correct rate around reversal* was calculated for the 15 trials preceding and 15 trials following each reversal and averaged across reversals within a session. Sessions in which no reversal was achieved were excluded from this analysis. Premature responses were excluded.
4. *Reversal number* was defined as the number of contingency reversals completed during a session.
5. *Responses per reversal* were calculated as the total number of correct and incorrect panel responses divided by the number of reversals. Responses made after the last reversal were excluded. If no reversal was achieved during a session, the total number of responses was used as the value.
6. *Premature responses* were defined as touches to the left or right panel before trial initiation by touching the center panel.

For the last day of DRL training and each PRL task, feedback-dependent choice strategies were additionally quantified using win-stay and lose-shift probabilities:

1. *Win-stay probability* was defined as the percentage of rewarded trials after which the mouse selected the same panel on the subsequent trial: Win-stay (%) = rewarded trials followed by the same choice / rewarded trials with a subsequent trial × 100.
2. *Lose-shift probability* was defined as the percentage of punished trials after which the mouse selected the opposite panel on the subsequent trial: Lose-shift (%) = punished trials followed by the opposite choice / punished trials with a subsequent trial × 100.

Premature responses were excluded from these calculations. The final trial of each session was also excluded because no subsequent choice was available to determine stay or shift behavior.

### Computational modeling

#### Reinforcement-learning model

To characterize the decision-making strategies used by mice during DRL and PRL tasks, we fitted reinforcement-learning (RL) models adapted from Hattori et al.^12^. The models were based on a modified Rescorla–Wagner framework in which the estimated values of the two choices were updated trial by trial according to the choice and its outcome. As in Hattori et al., the full model incorporated separate learning rates for rewarded and unrewarded outcomes, forgetting of the unchosen option, a constant choice bias, and sensitivity to the difference in value between the two options (**Figure 3A**).

The reward outcome on trial t, denoted R(t), was coded as 1 for a rewarded trial and 0 for an unrewarded trial. The values of the left and right options (Q_L_ and Q_R)_ were initialized to zero.

Following each trial, the value of the chosen option was updated according to:

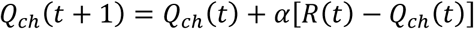

where separate learning rates were applied depending on trial outcome

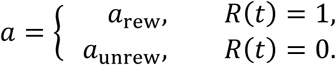

Thus, α_rew_ represents the extent to which a rewarded outcome increases the estimated value of the chosen option, whereas α_unrew_ represents the extent to which an unrewarded outcome decreases its estimated value. Both learning rates were constrained between 0 and 1.

When forgetting rate was included in the model, the value of the unchosen option was updated according to

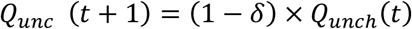

where δ is the forgetting rate and was constrained between 0 and 1.

The left choice probability on trial *t* was estimated from the difference between the left and right action values using the softmax choice rule described by Hattori et al.:

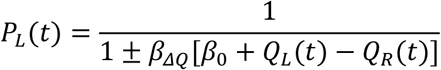

with *P_R_*_(_*t*) = 1 – *P_L_*_(_*t*).

Here, β_ΔQ_ represents sensitivity to the difference between the estimated values of the two options. β_0_ represents a constant bias toward one of the two options.

Five candidate models containing different combinations of these parameters were compared (**Table 1**). Model 1 contained only the constant bias term (β_0)_ and served as a baseline model without value updating. Model 2 contained separate learning rates (α_rew a_nd α_unr)_ and value sensitivity (β_ΔQ)_. Model 3 additionally included the constant choice bias (β_0)_, whereas Model 4 instead incorporated forgetting of the unchosen option (δ). Model 5 was the full five-parameter model and included α_rew_, α_unr_, β_ΔQ_, β_0_, and δ.

Model selection was first performed using WT mice. Five candidate RL models were compared using the first 80 trials from the final DRL session and the PRL 80–20 and PRL 70–30 tasks. Only completed left- or right-panel choices were included in the modeling. Premature responses were excluded. Model parameters were estimated for each animal by maximizing the likelihood of the observed trial-by-trial choices. Model fit was compared using the Akaike information Criterion (AIC), which accounts for both goodness of fit and the number of free parameters, with lower AIC values indicating better model fit. Models 4 and 5 yielded similarly low AIC values (**Figure 3B**). Model 5, which represents the full five-parameter model, was used for subsequent genotype comparisons.

#### Logistic-history model

As a complementary analysis that did not assume trial-by-trial updating of latent action values, choice behavior was also fitted using a logistic-history model based on Hattori et al.^12^. The left choice probability was modeled from rewarded-choice, unrewarded-choice, and outcome-independent choice histories over the preceding 10 trials:

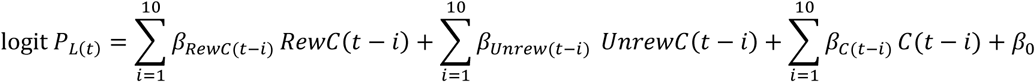

Rewarded-choice history was coded as 1 for a rewarded left choice, −1 for a rewarded right choice, and 0 otherwise. Unrewarded-choice history was coded analogously, whereas outcome-independent choice history was coded as 1 for a left choice and −1 for a right choice. These definitions follow the history predictors used by Hattori et al. The model therefore contained 30 history coefficients and one intercept (31 parameters in total).

The logistic-history model was fitted using L1-regularized logistic regression in glmnet, with regularization strength selected by ten-fold cross-validation using the one-standard-error rule. Because the model incorporated choice history over the preceding 10 trials, predictions began from trial 11 of each session, with the first 10 trials used to establish choice history. Choice history was reset between sessions. Prediction accuracy was assessed within each animal using two-fold cross-validation. The predictive performance of Model 5 was compared with that of the 31-parameter logistic-history model and a constant-bias baseline model.

#### Parameter-sweep simulations

To examine how individual reinforcement-learning parameters influence task performance, we performed Monte Carlo simulations using the fitted Model 5 parameters from WT mice. For each mouse, one parameter was systematically varied across 20 values while the remaining four parameters were fixed at that mouse’s fitted estimates. For each parameter value, 100 simulations of 80 trials were performed under each of four reward contingencies corresponding to the DRL 100–0 and PRL 80–20, 70–30, and 60–40 tasks. Reversal occurred when at least 12 of the preceding 15 choices were directed toward the currently correct option, with a minimum of 15 trials between reversals. Correct rate, number of reversals, and responses per reversal were calculated for each simulation. The 100 simulations were averaged within each mouse, parameter value, and task before statistical analysis.

### Statistical analysis

Behavioral data were analyzed using R and GraphPad Prism. Statistical tests were selected according to the experimental design and are specified in figure legends. Linear mixed-effects models were used for analyses involving repeated measurements, with mouse identity included as a random effect. Correct rate around reversals was analyzed using two-way repeated-measures ANOVA or, when data were missing, a linear mixed-effects model. One-way ANOVA or the nonparametric Kruskal–Wallis test was used for comparisons among multiple groups, with post hoc multiple comparisons performed as specified in the figure legends. The Mann–Whitney test was used for comparisons between two independent groups. Differences in the proportion of mice reaching task criteria over time were assessed using Kaplan–Meier analysis and the log-rank test. Associations between behavioral measures were assessed using Pearson’s correlation coefficient or linear mixed-effects models when repeated observations from individual mice were included. Normality was assessed using the Shapiro–Wilk test.

Reinforcement-learning models were implemented using custom MATLAB scripts, and model fits were compared using the Akaike Information Criterion (AIC). All statistical tests were two-sided unless otherwise specified, and *p* < 0.05 was considered statistically significant. Data are presented as mean ± SEM unless otherwise indicated.

## Acknowledgement

We thank Dr. Shintaro Otsuka and all the lab members for discussions on the manuscript.

## Author contributions

Conceptualization: AT, TM & AI. Data acquisition: JH, KP, TN, KH & AI. Data analysis: JH, KP, TN, TM & AI. Manuscript writing: JH, KP, TK, TM & AI.

## Data and code availability

All behavioral data is available upon request. Codes can be accessed from the following link: https://neurodata.riken.jp/r/Ito-Ishida/2026_Hilber_Adaptive%20decision-making%20in%20NDD%20models/ (PW: NDDflexibility)

